# Investigating mouse motor coordination using quantitative trait locus analysis to model the genetic underpinnings of developmental coordination disorder

**DOI:** 10.1101/2022.06.07.495138

**Authors:** Kamaldeep Gill, Jeffy Rajan Soundara Rajan, Eric Chow, David G. Ashbrook, Robert W. Williams, Jill G. Zwicker, Daniel Goldowitz

## Abstract

The fundamental skills for motor coordination and motor control emerge through development, from infancy to late childhood years. Neurodevelopmental disorders such as Developmental Coordination Disorder (DCD) lead to impaired acquisition of motor skills. This study investigated motor behaviors that reflect the core symptoms of human DCD through the use of BXD recombinant inbred lines of mice that are known to have divergent phenotypes in many behavioral traits, including motor activity. We sought to correlate behavior in basic motor control tasks with the known genotypes of these reference populations of mice using quantitative trait locus (QTL) mapping. We used twelve BXD lines with an average of 16 mice per group to assess the onset of reflexes during the early neonatal stage of life and differences in motor coordination using the open field, rotarod, and gait analyses during the adolescent/young adulthood period. Results indicated significant variability between lines in as to when neonatal reflexes appeared as well as significant line differences for all measures of motor coordination. Five lines (BXD15, BXD27, BXD28, BXD75, and BXD86) struggled with sensorimotor coordination as seen in gait analysis, rotarod, and open field, similar to human presentation of DCD. We identified three significant quantitative trait loci for gait on proximal Chr 3, Chr 4 and distal Chr 6. Based on expression, function, and polymorphism within the mapped QTL intervals, 7 candidate genes *(Gpr63, Spata5, Trpc3, Cntn6, Chl1, Grm7* and *Ogg1)* emerged. This study offers new insights into mouse motor behavior which promises to be a first murine model to explore the genetics and neural correlates of DCD.

## INTRODUCTION

The fundamental skills for motor coordination and motor control largely emerge through development, from infancy to late childhood to adulthood.^1^ When this developmental progression is perturbed as a result of rare genetic mutations, neurodevelopmental disorders can emerge.^1^ Neurodevelopmental disorders can cause atypical cognitive functioning, intellectual impairment, and/or motor developmental delays.^1^

Developmental Coordination Disorder (DCD) is a neurodevelopmental disorder affecting 5-6% of school-aged children.^2^ DCD is characterized by deficits in postural control, sensorimotor coordination, and motor learning that significantly affect a child’s ability to carry out activities of daily living, such as tying shoelaces, getting dressed, printing, or riding a bicycle.^2, 3^ Motor performance of children with DCD is more variable, less accurate, and slower than typically developing children.^2^ Children with DCD typically do not outgrow these motor impairments, rather they persist into adolescence and adulthood.^4^

DCD is a multifactorial disorder with unclear etiology, partly due to its heterogeneous nature. Research to date has suggested that genetics, family history, and altered brain development play a role in the phenotypic variability seen in DCD. Mouse models are a commonly used experimental platform to explore such heterogeneous human disorders in order to study the genetic, molecular, and behavioral aspects of human diseases.^5^ In modeling a complex disorder like DCD in the mouse, it is necessary to examine DCD-relevant behaviors across a spectrum of variation. The use of reference panels of mice, such as recombinant inbred (RI) lines, has proven to be a powerful approach to our understanding of a broad spectrum of disorders.^6^ The BXD family, currently the largest and best characterized mouse reference population, is composed of ∼150 lines.^7^ Additionally, the behavioral repertoire of BXD RI mice is diverse and reflects the substantial differences between the progenitor lines of the BXD lines, C57BL/6J and DBA/2J mice.^8^

To capture the motor variability seen in BXD RI mice, we devised a battery of motor behavior tasks (Figure 1) to mimic the motor deficits observed in human DCD (see Gill et al.^9^ for details). Using this approach, we can begin to identify candidate regions of chromosomes that harbor genes that underlie the behaviors of interest in DCD using Quantitative Trait Loci (QTL) analysis. In this paper, we focus on the variation of motor coordination phenotypes and corresponding QTLs in BXD lines of mice and assess whether we have access to variations in phenotypes that are relevant to DCD and correlations with genotypes.

**Figure 1.**
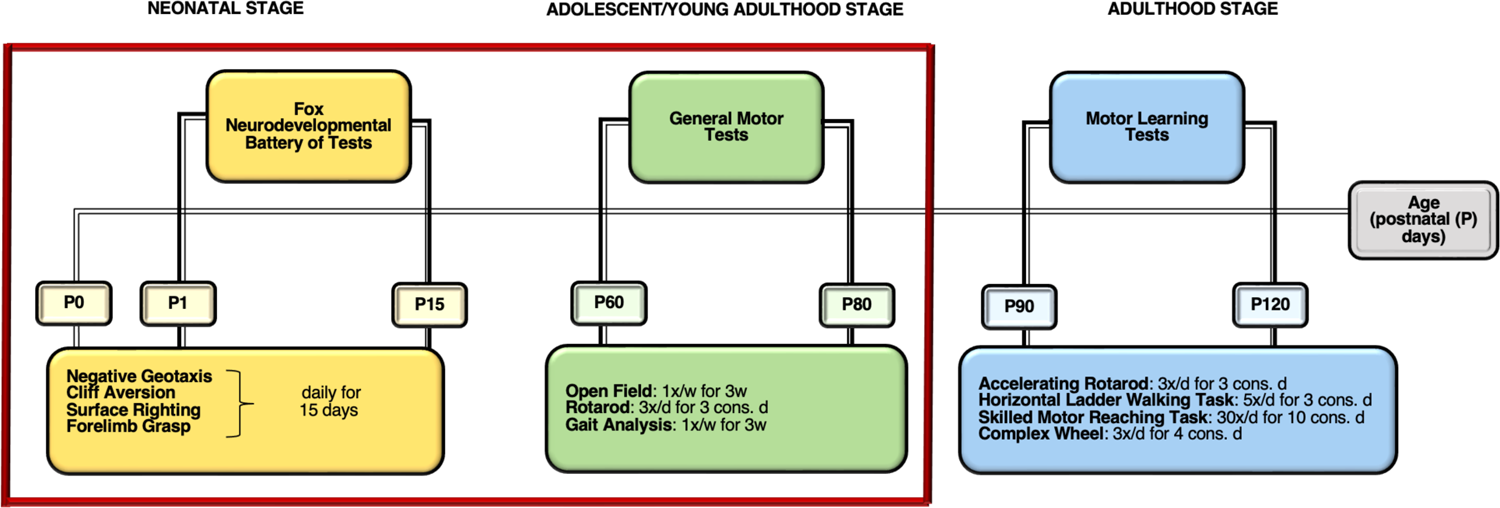
Workflow of the behavioral testing in Postnatal Day (P) P1-120. All three phases of testing are proposed based on the DCD-like behavior. [d-days, w-week, cons.-consecutive]

## METHODS

### Animals

The BXD breeding colony used to generate mice for this study was established from mice purchased from Jackson Laboratory (Bar Harbor, ME) and housed at the Transgenic Animal Facility, Center for Molecular Medicine and Therapeutics, University of British Columbia. A total of 280 mice from C57BL/6J (B6), DBA/2J (DBA), and 12 BXD lines (BXD1, BXD15, BXD27, BXD28, BXD32, BXD40, BXD45, BXD65a, BXD69, BXD75, BXD81, BXD86) aged from postnatal day 1 (P1)-P120, were tested in this study and housed in same sex groups of 2-5 at a temperature of 22.5°C, with a 12:12h light-dark cycle (lights on at 6:00am). Food and water were freely available, and bi-weekly cage changes took place at the end of the week. All testing took place during the morning hours (light phase) between 8:00-12:00 by co-authors KG and JR. All experiments were conducted in accordance with the guidelines of the Canadian Council of Animal Care, and all protocols were approved by the University of British Columbia Animal Care Committee (ACC).

The selection of each BXD strain to be examined was based on documented behavioral phenotypes and structural brain differences in the BXD population. Common assays, such as rotarod, open field, dowel test, and MRI findings were used to select strains that demonstrated core phenotypes that sat at the extremes and the mid-point for balance (latency to fall on rotarod, dowel test), motor coordination (spontaneous locomotion in open field), and cerebellar volume as measured by MRI. For example, strains were selected based on behavioral performance on motor coordination and learning tasks that ranged from poor to very good, in relation to human DCD phenotypes (Table 1).^8^ Further, BXD69, BXD75, BXD81, and BXD86 were selected for their performance variability ranging from poor to very good on the rotarod.^10^ Other strains, BXD32, BXD45, and BXD65a, were selected either due to observed variation in balance abilities on the dowel test and/or spontaneous locomotion in the open field.^8, 10^ BXD 1, BXD15, BXD27, BXD28, and BXD40 were selected due to their variation in cerebellar volume.^9^

**Table 1.** Selection of BXD strains based on motor-related traits

| GeneNetwork ID | Phenotype | Strain(s) |
| --- | --- | --- |
| 10921 | Cerebellar volume | BxD1/TyJ<br>BxD15/TyJ<br>BxD27/TyJ |
| 10921; 11819 | Cerebellar volume; dowel test | BxD28/TyJ |
| 11819 | Dowel test | BxD32/TyJ |
| 10921 | Cerebellar volume | BxD40/TyJ |
| 11014 | Open field behavior | BxD45/RwwJ<br>BxD65a/RwwJ |
| 11004 | Rotarod performance | BxD69/RwwJ<br>BxD75/RwwJ |
| 11014; 11005 | Open field behavior; improvement in rotarod | BxD81/RwwJ |
| 11004; 11819 | Rotarod performance; dowel test | BxD86/RwwJ |

### Procedures: Behavioral Testing

Detailed information on the study timeline is shown in Figure 1 and rationale and procedural details for the behavioral tasks are outlined in Gill et al.^9^ This paper specifically focuses on the neonatal and the adolescent/young adulthood stage in BXD mice. During the neonatal stage, a battery of reflexes was studied for each pup, daily for the first 15 days of life to examine postnatal maturation of nervous system and motor behavior (Figure 1).^11^ Specifically, we examined negative geotaxis, cliff aversion, surface righting, and forelimb grasp in neonate mice. In the adolescent/young adult stage of life, we measured motor coordination using open field (total distance traveled, amount of time spent in the center versus the periphery, time spent moving versus not moving, and velocity), standard rotarod (latency to fall), and gait analysis (body speed, leg combination, duty factor, step cycle, swing duration, stance duration, and posterior exterior position) between the ages of P60-P81 (Figure 1). The order of testing for all motor tasks was randomized for each litter on each testing session to limit the test order interaction and the effect of fatigue on performance.

### Behavioral Data Analysis

<u>Statistical analysis</u> was carried out using IBM SPSS Statistics Version 25.0. Data were analyzed separately for each test by one-way ANCOVA using strain as an independent variable and body weight and sex as covariates. Post hoc comparisons were performed when appropriate using the Bonferroni method. To control for potential litter effects, we used the strain averages for each of the respective variables. Further, using strain average increases the power to detect the genetic effects.^12^

An average of all trials per day for each task was taken for each of the testing days. The effect of the strain over the several testing days was explored using a two-way ANCOVA with body weight and sex as covariates. To examine motor performance improvements, a difference score was calculated (mean of day 3 or 4 - mean of day 1). All measures are reported as the mean +/- S.E.M. and the level of significance reported for all comparisons is p< 0.05. <u>Heritability</u> is the proportion of phenotypic variance that is explained by genetic differences. In fully homozygous strains, such as the BXD, additive genetic variance (Va) is the main source of differences between strains, and our estimates can be considered close to what is typically called narrow-sense heritability (h^2^). h^2^ was estimated as the fraction of variance explained by strains in a simple ANOVA model.

### QTL mapping

The behavioral trait data were uploaded into GeneNetwork (www.genenetwork.org), an open-access database which contains genotypes for the BXD family and phenotypic information that was entered by the study team. For each phenotype, we mapped QTLs using the fast linear regression equations of Haley and Knott. ^86^ The likelihood ratio statistic (LRS) was computed to assess the strength of genotype–phenotype associations. When appropriate, outlier values for specific traits were winsorized. A test of 2000 permutations was performed to evaluate the statistical significance of associations, and identify suggestive (p < 0.63) and significant (p < 0.05) QTLs. The bootstrap test was implemented to estimate the confidence interval by generating multiple bootstrap datasets by randomly selecting lines with substitution from the original RI set. Confidence intervals around the significant LRS score peaks were calculated using 1.5 logarithm of the odds confidence interval.

### Candidate gene analysis

Using the online tools in GeneNetwork, the genes within the QTL confidence interval were evaluated based on three criteria: (1) gene expression within brain and skeletal muscle region of interests; (2) relevant gene functions from previous literature; and (3) the presence of polymorphisms (i.e., non-synonymous SNPs) as the phenotypic difference is assumed to be associated by genetic variation and considered as good candidate genes. The expression, functional, and phenotypic information for each of the genes located within the significant QTLs^13, 14^ were assessed using Allen Brain Atlas, Mouse Genome Informatics (MGI) database, and PubMed. Secondly, literature searches were conducted to determine if each gene under the significant QTL peak had a previously reported role in motor skills and learning. Lastly, GeneNetwork variant browser was used to identify SNP variants specifically nonsynonymous polymorphism between parental haplotypes.

## RESULTS

### Neurodevelopmental Battery

The offspring of all strains exhibited sensorimotor reflexes within the average timeline as specified by Fox^11^ and Heyser et al.,^15^ indicating that all strains of mice had typical neurological development. Although all strains displayed sensorimotor reflexes within the typical age range, there were significant strain differences, measured by average performance across the 15 days of testing for each strain: negative geotaxis [F (13, 274) = 11.154, p<0.001], cliff aversion [F (13, 274) = 12.785, p<0.001], surface righting [F (13, 274) = 8.805, p<0.001], and forelimb grasp [F (13, 274) = 5.532, p<0.001] (Figure 2; Table 2). For negative geotaxis, a significant difference was observed between B6 and DBA parental strains. Similarly, in cliff aversion, each strain displayed variability in the time taken to turn away from the edge to the secure side. These differences persisted for the entire 15 days of testing, with B6 and BXD86 having a statistically significant difference in time to observe the reflex. Contrary to negative geotaxis and cliff aversion, surface righting showed less variability in time taken to restore a pup’s position from dorsal to its normal prone position. In forelimb grasp, strains displayed less variability in time to hold the rod and remain suspended above platform, with no significant difference between strains.

**Figure 2.**
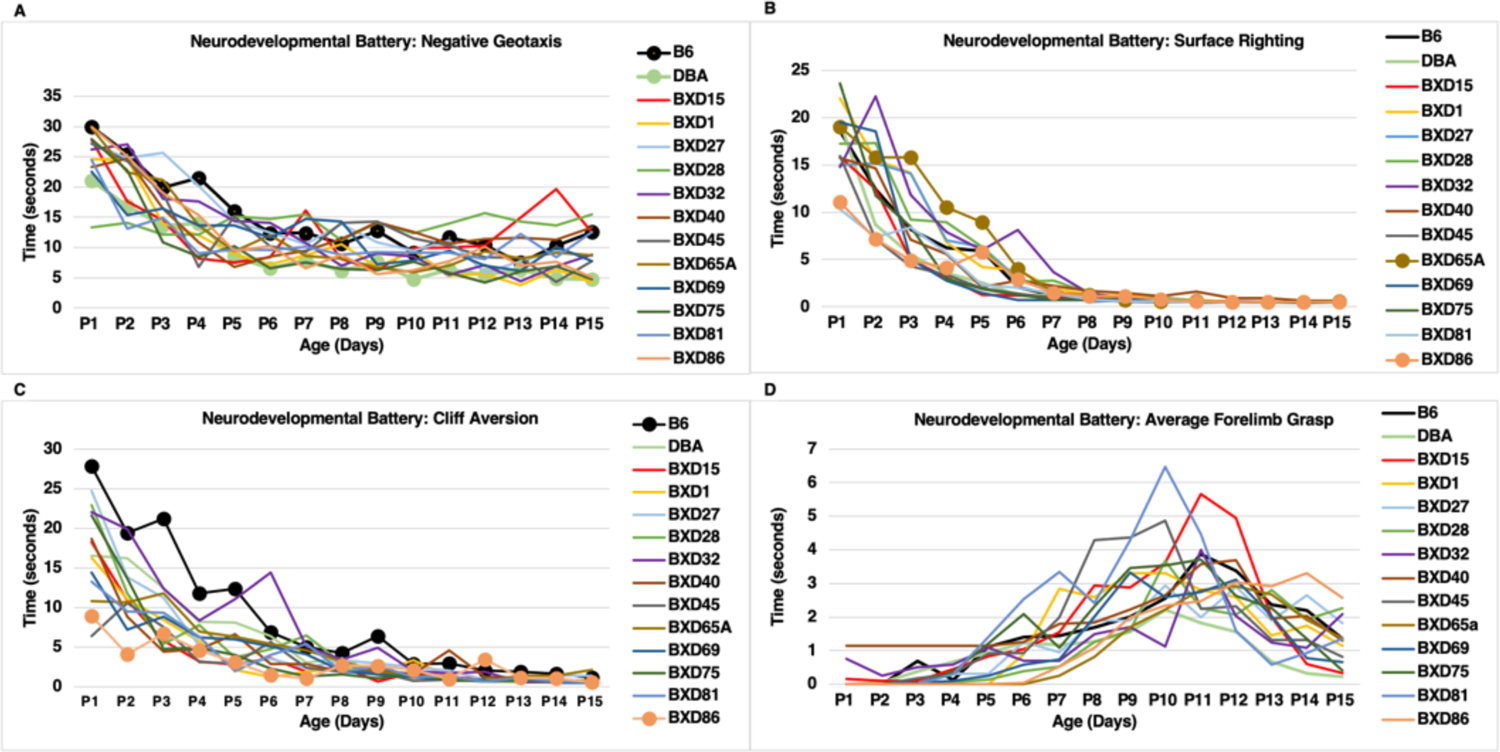
Graphs illustrate strain response time on Fox neurodevelopmental battery over the first 15 postnatal days. (A) negative geotaxis; (B) cliff aversion; (C) surface righting; and (D) forelimb grasp. The wide range of colors shows individual strain performance.

**Table 2.** Summary of behavioral tests and statistical analysis

| Test | Phenotype | Parameter | ANCOVA (all $p < 0.001$) |
| --- | --- | --- | --- |
| Fox Neurodevelopmental Battery | Nervous system maturation | Negative geotaxis | $F(13, 274) = 11.15$ |
| | | Cliff aversion | $F(13, 274) = 12.79$ |
| | | Surface righting | $F(13, 274) = 8.81$ |
| | | Forelimb grasp | $F(13, 274) = 5.53$ |
| Open field | Locomotor activity | Total distance travelled | $F(13, 280) = 4.54$ |
| | | Time spent in the center | $F(13, 280) = 10.54$ |
| | | Time spent in the periphery | $F(13, 269) = 34.80$ |
| | | Time spent moving | $F(13, 274) = 15.25$ |
| | | Time spent not moving | $F(13, 280) = 20.30$ |
| | | Velocity | $F(13, 280) = 3.47$ |
| Rotarod | Motor coordination, balance | Baseline performance | $F(13, 280) = 5.38$ |
| | | Performance improvement | $F(13, 280) = 4.83$ |
| Gait analysis | Motor coordination,<br>postural control | Body speed | F (13, 236) = 10.12 |
|  |  | Leg combination | F (13, 236) = 15.00 |
|  |  | Stance duration | F (13, 236) = 23.54 |
|  |  | Swing duration | F (13, 236) = 15.74 |
|  |  | Posterior extreme position | F (13, 236) = 10.74 |
|  |  | Step cycle | F (13, 236) = 23.54 |
|  |  | Duty factor | F (13, 236) = 13.93 |

### Motor Activity

#### Open Field

An open field test was performed as an indicator of general locomotor activity. The total distance traveled, amount of time spent in the center versus the periphery, time spent moving versus not moving, and velocity were analyzed (Figure 3). There were significant strain differences for total distance travelled, time spent in the center, time spent not moving, and velocity over the 10 minute testing period (Table 2). Figure 3 shows that BXD27, BXD28, and BXD32 spent the least amount of time moving. BXD27, BXD69, BXD75, BXD81, and BXD86 had the lowest velocity of movement. When considering the distance traveled within the 10 minutes of testing, three of these four strains, BXD27, BXD69, and BXD75, also logged the least distance. We did not find any sex difference or significant correlations between weight and open field parameters (p>0.05).

**Figure 3.**
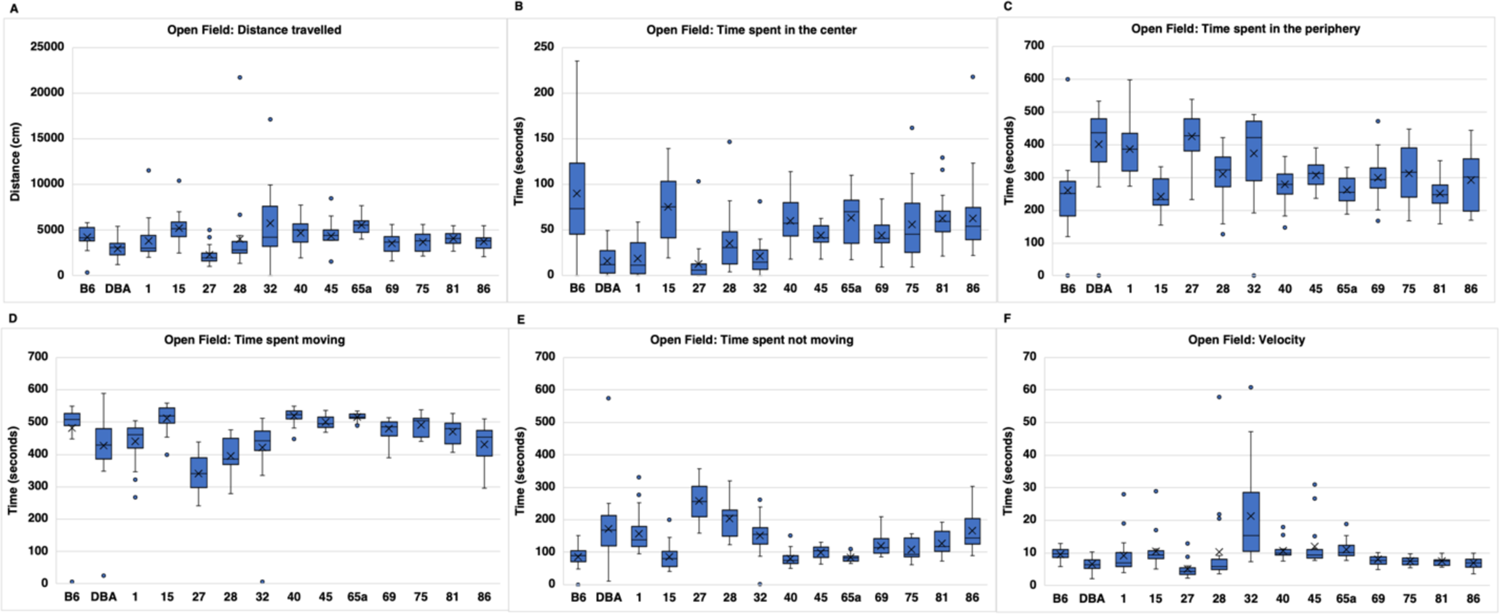
Graphs illustrating strain performance in open field parameters: (A) distance travelled; (B) time spent in the center; (C) time spent in the periphery; (D) time spent moving; (E) time spent immobile; and (F) velocity. The x in the box depicts the mean and the bottom line of the box depicts the median.

### Motor Coordination

#### Rotarod

To investigate motor coordination in our BXD panel, the rotarod test was performed using a constant speed task. We determined rotarod performance by measuring the latency to fall from the rotarod. A one-way ANCOVA revealed a significant effect of strain on rotarod performance, particularly on the first day of testing (Table 2). Specifically, BXD28 BXD32, BXD65a, BXD69, BXD75, and BXD86 performed below average on the first day of testing; BXD69 and BXD86 had the shortest latency to fall (Figure 4). Subsequently, a two-way ANCOVA showed a significant interaction between BXD strain and day of testing.

**Figure 4.**
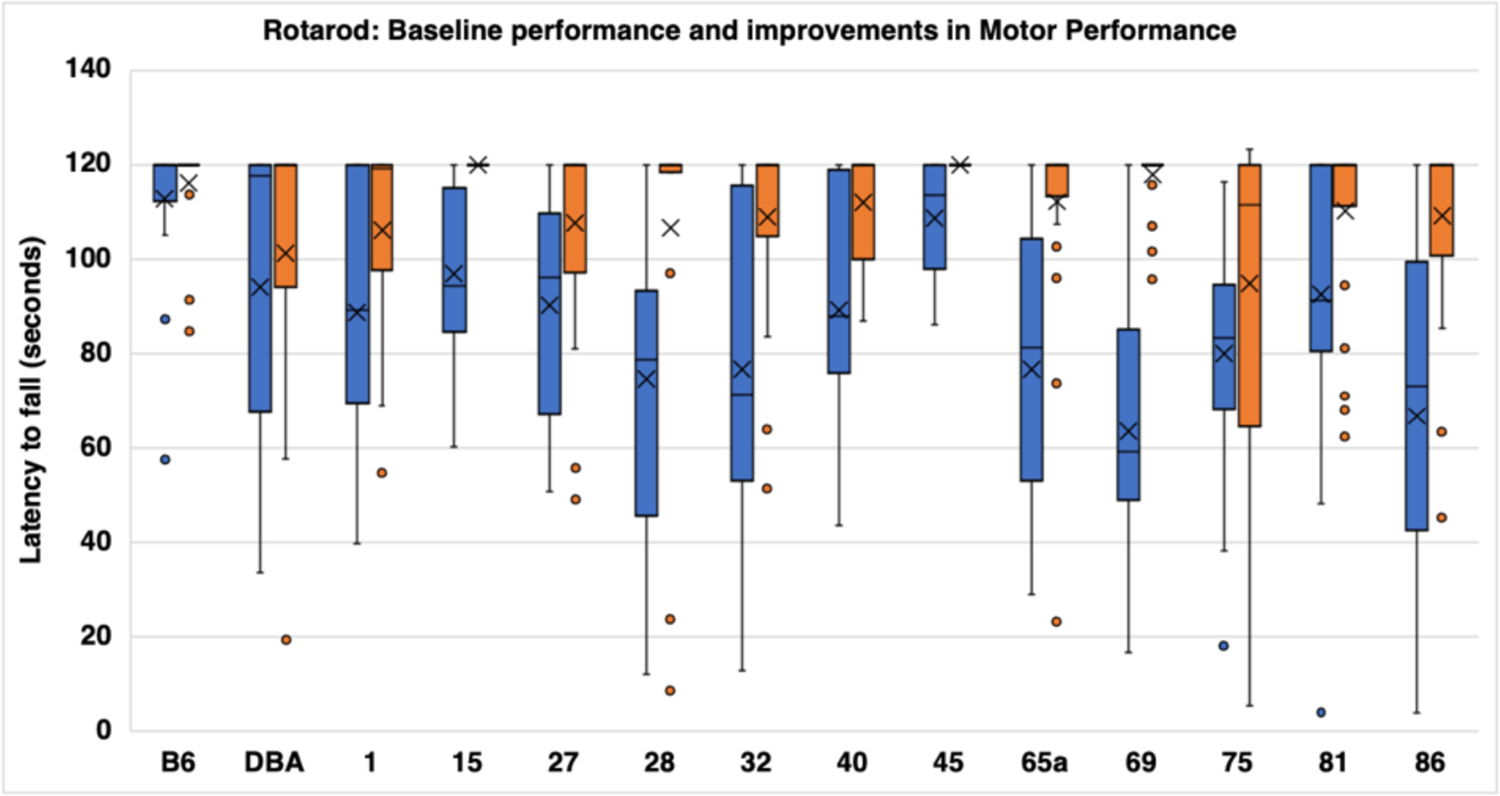
Strain performance improvement on rotarod. Blue represents day 1 strain performance in latency to fall, Orange represents day 3 strain performance in latency to fall. The x in the box depicts the mean and the bottom line of the box depicts the median.

To calculate motor performance improvements, the rotarod performance on Time 3 was subtracted from the performance on Time 1 (mean of Time 3 - mean of Time 1). This analysis was directed to determine if previous experience and practice from Times 1 and 2 influenced performance on the last day of testing, indicating motor improvements if the mice became more adept on the rotarod. There were main effects of strain on motor learning (Table 2). Figure 4 illustrates the performance improvements from Time 1 to Time 3. We find that BXD27 and BXD 28, along with the parental DBA strain, show the lowest relative improvement. There was no correlation between adult body weight and motor learning and no sex differences were observed on rotarod performance (p>0.05).

#### Gait Analysis

To evaluate motor coordination, specifically postural control, gait analysis was performed. Gait performance was determined by measuring body speed, leg combination, duty factor (i.e., proportion of the step cycle where the leg is in contact with the ground), step cycle, swing duration, stance duration, and posterior exterior position (i.e., position of the leg relative to the end of stance phase) (Figure 5). To detect between-strain differences in gait parameters, an ANCOVA was performed. Statistically significant differences among the strains were identified for all of the gait parameters: body speed, leg combination, duty factor, step cycle, swing duration, stance duration, and posterior exterior position (Table 2). Post hoc tests for all gait parameters revealed that BXD15, BXD27, and BXD86 displayed the most variable performance (Figure 5). In particular, these strains had lower body speed and stance duration with a longer step cycle and duty factor.

**Figure 5.**
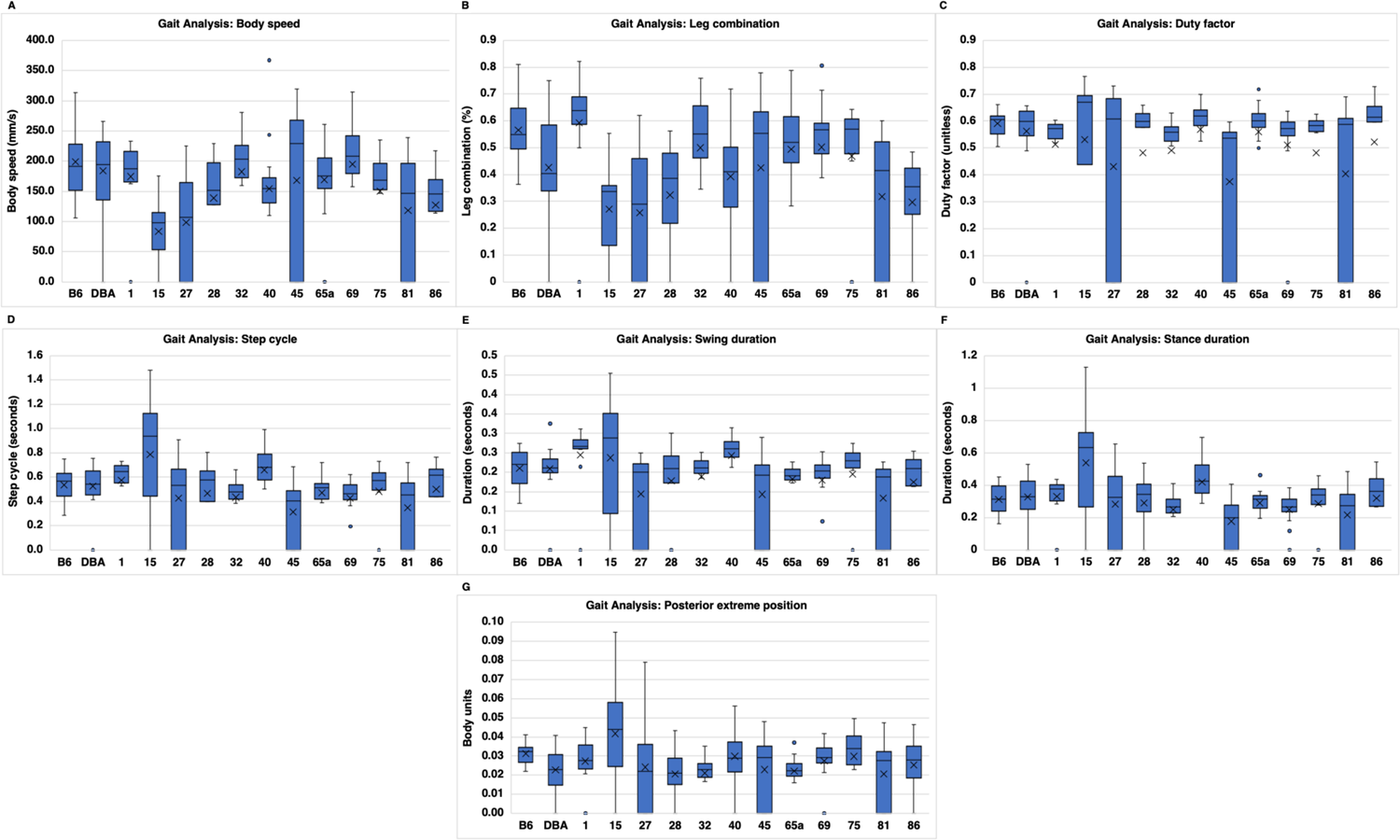
Graphs illustrating strain performance in gait parameters. (A) body speed; (B) leg combination; (C) duty factor; (D) step cycle; (E) swing duration; (F) stance duration; and (G) posterior extreme position. The x in the box depicts the mean and the bottom line of the box depicts the median.

### Heritability

For each of 35 phenotypes, heritability (h^2^) was calculated. Phenotypes with high heritability have a large effect of genometype on the phenotypic variation. Body weight, as expected, was highly heritable, at h^2^ 0.43 – 0.55, providing reassurance of correct strain assignment. We had greater power to identify QTL for traits with high heritability, since power to detect QTL is dependent upon heritability, locus effect size, number of genometypes examined, and number of biological replicates. We therefore expected to be more likely to detect QTL for phenotypes such as step cycle (h^2^ = 0.57), stance duration (h^2^ = 0.57), time not moving (h^2^ = 0.49), swing duration (h^2^ = 0.46), leg combination (h^2^ = 0.45) and duty factor (h^2^ = 0.43). We note that four of these five are gait phenotypes, and therefore that aspects of gait are strongly influenced by genometype in the BXD family. At least some of this shared heritability may be due to high correlations between traits (e.g. correlations between step cycle, swing duration and stance duration are r^2^ 0.56 – 0.96), which would likely result in shared QTL positions.

### QTL Analyses

A summary of the QTLs is presented below and in Table 3. We found three significant QTLs, all from traits associated with gait analysis. A total of 13 suggestive QTLs were identified that spanned all of the behavioural tests given.

**Table 3.**
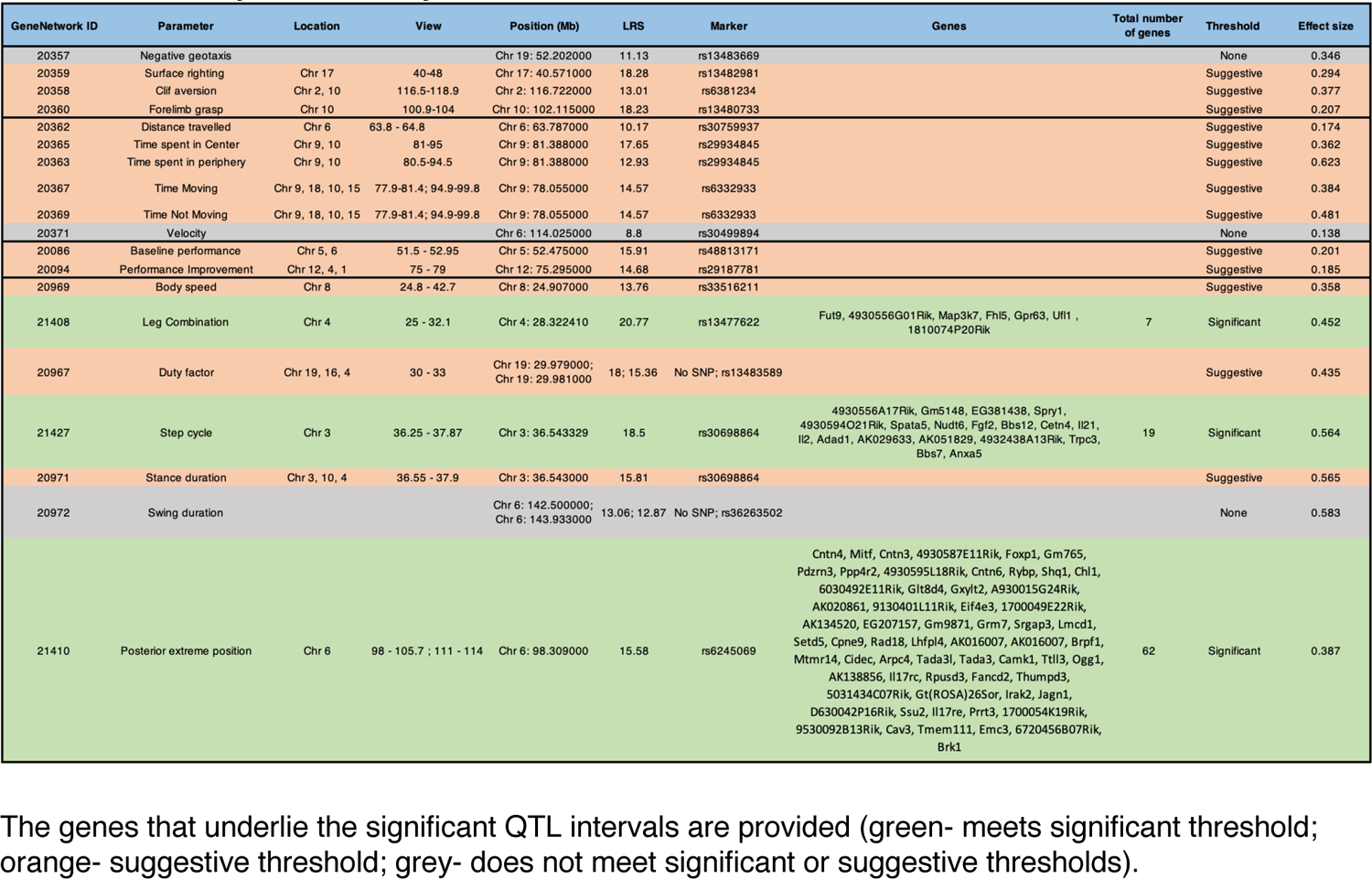
Summary of QTL analysis of behavioral traits

#### Neurodevelopmental Battery

The genome-wide QTL analysis did not identify any significant QTLs for any elements of the battery but suggestive QTLs were noted for righting reflex on Chr 17; for cliff aversion on Chr 2 and Chr 10; and for forelimb grasp on Chr 10 (Figure S1).

#### Open Field

Genome-wide QTL mapping for quantified traits in the open field test did not identify any significant QTLs. Several suggestive QTLs were found on Chr 9, Chr 18, Chr 10 and Chr 15 for time spent moving and not moving; on Chr 10 and Chr 9 for time spent in center and periphery; and on Chr 6 for total distance traveled (Figure S2).

#### Rotarod

The genome-wide QTL map did not identify any significant QTLs for rotarod measures but identified suggestive QTLs for baseline performance on Chr 5 and Chr 6; and for performance improvement on Chr 12, Chr 4 and Chr 1 (Figure S3).

#### Gait Analysis

The genome-wide QTL map for gait analysis identified three significant QTLs in different gait parameters (Figure 6). For leg combination, a significant QTL was located to the proximal end of Chr 4 [GeneNetwork Trait ID_21408] and spanned from 25 to 32.1 Mb with a LRS score of 20.77. A significant QTL for step cycle was also mapped to the proximal end of Chr 3 and it spanned from 36.25 to 37.87 Mb with a LRS score of 18.5 [GeneNetwork Trait ID_21427]. A significant QTL for posterior extreme position was mapped to the distal end of Chr 6 [GeneNetwork Trait ID_21410] and spanned from 98 to 105.7 and 111 to 114 Mb with a LRS score of 15.58. Other parameters had suggestive QTLs: body speed on Chr 8; duty factor on Chr 19, Chr 16 and Chr 4; stance duration on Chr 3, Chr 10 and Chr 4; leg combination on Chr 5, 16 and 19; and duty factor on Chr 10, 4, 16 and 19 (Figure S4).

**Figure 6.**
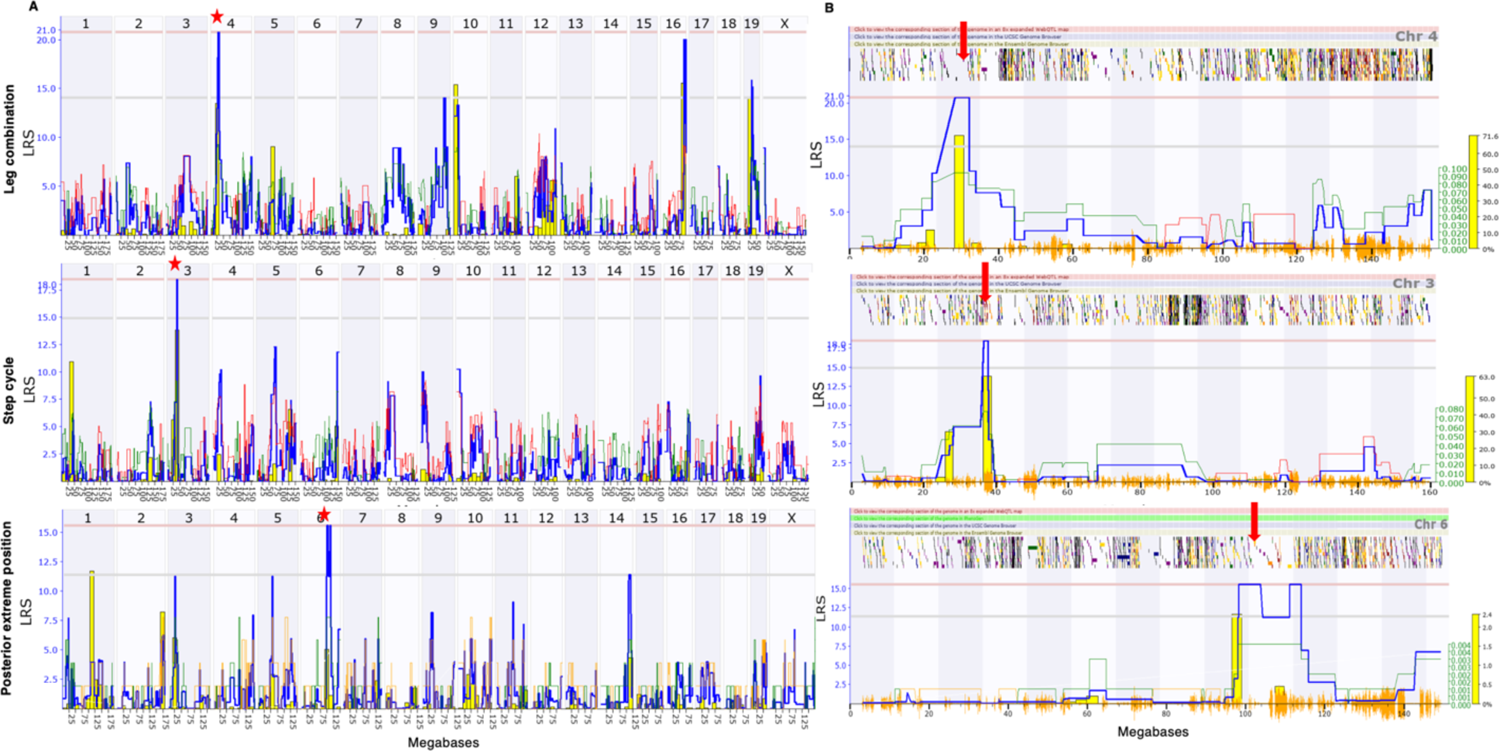
Genome-wide linkage map of leg combination, step cycle and posterior extreme position (top to bottom) of gait analysis to determine postural control. The overall blue trace shows the LRS. (A) Genome-wide QTL map showing a significant QTL on chromosome 4 for leg combination; chromosome 3 for stance duration annd chromosome 6 for posterior extreme position (B) Interval QTL map of chromosome 4, 3 and 6 using three test week performance with bootstrap analysis. The lower gray horizontal line represents suggestive LRS genome-wide threshold at p ≤ 0.63. The upper pink horizontal line represents significant LRS genome-wide threshold at p ≤ 0.05. The bottom orange marks indicate SNP density. [asterisk (*) indicates significant QTL; down arrow (↓) indicates significant QTL interval with genes].

#### Candidate Genes for Significant QTLs

With the identification of significant QTLs for three traits in gait analysis, our attention focused on the best candidate genes that might underlie each QTL. To this end, we identified 88 genes that were mapped to the three significant QTL regions (Table 3).

As described in the Materials and Methods section, a series of criteria were employed to prioritize promising candidates based upon: (1) the gene is expressed in central nervous system (CNS) and/or skeletal muscle; (2) the gene is associated with a motor function related to DCD-like behavior; and (3) presence of nonsynonymous SNPs in coding region of the gene. The expressed sequence tags and Riken clones within the gene list were not evaluated since they are currently non-annotated in terms of expression pattern and function. Of all the genes observed across three significant QTLs, 59 genes met criterion 1; 7 of these 59 genes met criterion 1 and 2 (Table 3); and 3 of these 7 genes met all three criteria (Table S1).

Tissue-specific expression is an important factor for determining the role of genes in a given disorder. The motor coordination problems are significant for DCD diagnosis and are likely associated with genes involved in brain development and/or skeletal muscle function.^16^ Hence, the initial assessment of the 88 gene expression profile datasets were screened to determine expression in brain and skeletal muscle tissues relative to other tissues using Allen brain atlas, NCBI and MGI databases. Of the 88, we identified 59 genes that had higher expression in CNS and skeletal muscles (Table S1). Secondly, the functional role of 59 genes was explored using NCBI resource and identified the following genes: *Gpr63, Spata5, Trpc3, Cntn6, Chl1, Grm7 and Ogg1*as meeting criterion 2. Lastly, the SNP variant browser was used to scan the seven genes, above, and identified three genes that have non-synonymous polymorphisms. Thus, 3 candidate genes, *Spata5*, *Cntn6* and *Chl1,* within two significant QTL regions genes that met all three criteria (1,2 & 3) and are therefore considered as priority genes to explore in relation to the clinical presentation of DCD.

## DISCUSSION

The results of this study show that there is significant variability in posture, balance, motor activity, and sensorimotor coordination amongst the BXD RI strains as measured by gait analysis, rotarod, and open field behavioral assays and this bodes well for using the BXD strains to explore the structural, functional, and genetic aspects of motor behavior. To our knowledge, this is the first study to explore the motor variability in BXD strains of mice in order to explore DCD-like phenotypes and their genetic underpinnings. Of particular note is that strains of BXD mice share core symptoms displayed by individuals with DCD. For example, BXD75 and BXD86 had motor coordination difficulties that are not unlike individuals with DCD who have poor static and dynamic balance,^17–19^ decreased spontaneous locomotion^20–24^ and altered gait patterns.^5,25, 26^ As seen in BXD15, BXD27, BXD28, BXD75, and BXD86 strains of mice, children with DCD also struggle to improve their motor performance with increased opportunity to practice. Individuals with DCD continue to show variable and less accurate motor performance over time, which is also seen in our BXD strains of mice. The observed gait differences across BXD strains are also consistent with gait patterns of children with DCD, who show abnormal walking patterns,^81^ including bilateral asymmetry,^82–84^ decreased ability to coordinate lower limbs,^82, 84^ greater step width,^85^ and longer time in stance phase.^85^ Thus, the behavioral phenotypes seen in this study are similar to what may be seen in individuals with DCD and have the potential to offer insight into the genetic and neuroanatomical underpinnings of motor coordination impairments. At first blush, these results show great potential for a murine model to study the etiology of DCD.

### Hypthesis Driven Selection of Genotypes and Phenotypes

Generally, testing a small number of strains yield fewer behavioral differences and a diminished probability for detecting QTLs; however, in this study, we saw significant inter-strain variability for all traits studied as well as the detection of significant QTLs. These findings would seem likely due to hypothesis-driven selection of BXD strains to study, along with having a population with large inherent differences as embodied in B6 vs DBA and the BXD strains that emerge from the recombinant inbred cross. Our results were undergirded by the extensive phenotyping that has been done in the field which provided many selection criteria to choose from. One selection criteria was based on the observed motor phenotypes in BXD strains of mice. As such, strains that showed variable performance on motor coordination tasks were chosen, along with those who had variation in brain structures that are associated with motor skills.

While some strains were selected based on known brain structural differences, others were selected based on the variability in motor performance in posture, balance, and locomotor activity tasks. Thus, BXD75 and BXD686 were amongst the strains that were chosen based on motor performance. Similar to previous findings, we also observed BXD75 and BXD86 to struggle with motor coordination and balance.^8, 10^ On the open field test, Brigman et al.^10^ and Philip et al.^8^ found that BXD75 and BXD86 presented with sedentary behavior and below average general locomotion, which may be due to motor coordination impairments. While the reason for difficulties in general locomotion and balance for both BXD75 and BXD86 may not be clear, other studies identify why these strains may struggle with motor coordination in general. Loos et al.^27^ and Laughlin et al.^28^ evaluated strain differences in attentional performance in a serial reaction time task and an operant learning task, respectively. Loos et al.^27^ found that BXD75 struggled with attentional performance; while Laughlin et al.^28^ found that BXD86 had difficulty with behavioral flexibility in order to change or stop a response.

It is known that attention and behavioral flexibility are needed for motor coordination of functional motor tasks.^27, 28^ Impaired attentional abilities and lack of ability to change or stop a response, may both depend on neuroanatomical differences. Jan et al.^29^ and Gaglani et al.^30^ found that BXD75 and BXD86 both have lower cerebral cortex volume, particularly somatosensory cortex volumes. The somatosensory cortex plays a critical role in processing afferent somatosensory input and contributes to the integration of sensory and motor signals that are necessary for skilled movement; abnormalities in this structure may lead to motor dysfunction.^29^ Therefore, it is plausible to suggest that structural differences in the somatosensory cortex may be contributing to attentional and behavioral inflexibility in BXD75 and BXD86 that lead to poor motor coordination and balance.

A second selection criterion was based upon neuroanatomical phenotypes which are important to assess in neurodevelopmental disorders, such as DCD, because anatomy is closely connected to behavior.^31^ As such, this presented another criterion for selection and our initial selection criteria applied to BXD strains had to do with varying cerebellar volume as this structure is intimately related to motor coordination. In fact, abnormalities in cerebellar structure and functions are hypothesized to be the main neural correlates implicated in many neurodevelopmental disorders, including DCD.^4^ Badea et al.^32^ compared various brain structures in BXD strains of mice and found that BXD27 had the lowest total cerebellar volume; whereas BXD15 and BXD28 had moderate cerebellar volume and BXD1 had the highest cerebellar volume, when compared to each other and to the wildtype parental strains. It is known that the cerebellum is involved in motor coordination and learning;^33^ and as such, we found that BXD15, BXD27, and BXD28, the strains with smaller cerebellar volumes show substantial alterations in motor coordination and balance as seen in the rotarod, open field, and gait analysis. On the other hand, BXD1 mice that present with the highest cerebellar volume, comparable to B6, performed well on all motor coordination tasks and presented with a typical gait pattern. Our findings are further strengthened by Ceccarelli et al.^34^ who studied the effects of altered cerebellar development on motor coordination and balance in mice. They found that gene knockout of Tsi21/Btg2 in the Btg family of mice caused altered migration of cerebellar granule precursors causing abnormal cerebellar volume and thickness, which led to motor coordination and balance impairments, as seen on the rotarod.^34^ Further, Galante et al.^35^ found that abnormal cerebellar structure, consisting of reduced cerebellar neuronal number and consequently reduced neuronal density in the cerebellum of Tc1 mice, led to major deficits in motor coordination and balance on the rotarod, decreased motor performance improvements on consecutive days of testing on the rotarod, reduced ability to habituate to the environment in the open field, and differences in gait patterns.^35^ Veloz et al.^36^ also found that mice with abnormal cerebellar structure that includes lack of Purkinje cell output exhibited short and irregular strides recorded by footprint analysis, had difficulties with keeping balance on the rotarod, and displayed reduced open field locomotion activity.^36^ A typical sign of cerebellar dysfunction is gait ataxia, which is characterized by balance problems and walking abnormalities, similar to what was observed for BXD15, BXD27, and BXD28.^36^

### Motor Behavior QTLs

Despite the limited number of BXD lines used in this analysis, we were able to detect 3 significant QTLs. This may be due to a number of features of the present work, including fortuitous outcomes, the so called Winner’s Curse.^88^ The pre-selection of lines for testing (see above) is likely to provide a more robust filter to enable stronger correlations between behavior and the genome – that is, strains at the poles of the distribution were chosen, producing a larger effect size. In addition, the large number of mice used in each group reduced environmental variation, allowing a more accurate estimate of strain mean for each phenotype, and therefore increased power to identify QTL. We can be slightly reassured by the fact that our most significant QTL also had high heritability, supporting the strong genetic effect on these traits.

The three genes, *Cntn6*, *Chl1*, and *Spata5*, that passed all three criterial proved to be very interesting candidates. *Cntn6 and Chl1* genes emerged from the posterior extreme position parameter. Genetic mutation of *Chl1* in mice have shown that partial or complete deletion leads to cognitive, behavioral, motor, and coordination deficits.^45, 46^ *Cntn6*, a member of the immunoglobulin superfamily is expressed exclusively nervous system development. Behavioral studies have shown that Cntn6-deficient mice displayed impaired motor coordination.^47^ We also identified *Spata5* from the step cycle gait parameter. Spata5 is spermatogenesis related gene and a member of the ATPase family. They are abundantly expressed throughout the brain and nutations of the gene in the human population have been found to result in major perturbations in brain development. ^39^ The details of Spata5 in various cellular processes, particularly in the brain development is still to be defined..

### Genes, gait behavior, and previous QTL analyses

There is an indication that DCD has an underlying genetic component due to its high heritability (∼70%).^52, 53^ Evidence suggests familial clustering of DCD,^54^ as well as an increased rate of copy number variations.^55^ To explore such gene-phenotype correlations in past work on motor control we examined previously acquires mouse phenotypic data in MGI. Here we find that the chromosomal location of the gait QTLs overlapped with abnormal phenotypes that included emotionality, stress^56^ and motor coordination,^41^ anxiety,^57^ and locomotor activity.^58^ These phenotypes map provocatively to many children with DCD who experience stress particularly in physical education classes.^59^ Sometimes, these children have difficulty with behavioural and emotional responses^60^ and show more aggression during play than their same-aged peers,^61^ including unprovoked hitting, kicking, and grabbing.^62^ Problems collectively experienced in the physical, cognitive, emotional, and social domains of development could limit the skills and resources from which children with DCD can draw to cope adaptively with stressful situations. Persistent problems experienced in this context will negatively impact children’s functioning and well-being.^63^ Overall, the investigation of previously reported studies within significant mapped QTL identified overlapping loci for other annotated abnormal phenotypes that may be associated with motor-related phenotypes.

The suggestive QTLs draw attention to loci that may be worth tracking and screening of rare variants. Suggestive QTL were mapped to various chromosomal regions and they overlapped with various phenotypes in MGI using the “phenotype, alleles, and disease models” query. Thus for gait parameters [(Chr 8: 24.8 to 42.7 Mb); (Chr 19: 30 to 33 Mb); (Chr 3: 36.55 to 37.9 Mb)] - body speed, duty factor, and stance duration - the query output generated various phenotypes including ethanol consumption,^64^ motor coordination^41, 47^ and learning and memory.^65^ For rotarod parameters [Chr 5 (51.5 to 52.95 Mb) and Chr 12 (75 to 79 Mb)] - baseline performance and performance improvement - the query output generated various phenotypes including weight control,^66^ motor coordination,^67^ locomotor activity,^68^ organ weights, and limb lengths.^69^ When we queried the suggestive QTLs for open field parameters - distance travelled, time spent in center, time spent in periphery, time moving and time not moving - [(Chr 6: 63.8 to 64.8 Mb); (Chr 9: 81 to 95 Mb); (Chr 9: 80.5 to 94.5 Mb); (Chr 9: 77.9 to 81.4 and 94.9 to 99.8 Mb)] the query output generated phenotypes that included locomotor activity,^68^ mood,^70^ learning and memory,^71^ cerebellar development^72^ and anxiety-like behavior.^73^ Finally, we inputted the suggestive QTL location for the neurodevelopmental battery [(Chr 15: 40 to 48 Mb); [(Chr 2: 116.5 to 118.9 Mb) and (Chr 10: 100.9 to 104 Mb)] the query output generated anxiety,^74^ body weight,^75^ locomotion activity,^76^ feeding,^77^ and circadian rhythms.^78^

In summary, the rich genetic and phenotypic diversity in the BXD RI strains and analysis of behavioral traits allowed us to identify three significant chromosomal regions that correlate with aspects of gait, which is known to be atypical in children with DCD in terms of timing, coordination, and symmetry. ^5, 25, 26^ Using bioinformatic resources, we identified 3 candidate genes in these QTL regions (*Cntn6*, *Chl1*, and *Spata5*). In addition, overlapping loci with abnormal phenotypes (e.g., anxiety, motor coordination, locomotion activity) were observed at some of the significant, mapped regions. The most promising candidate genes within the QTLs contain nonsynonymous sequence polymorphisms that may be involved in the regulation of motor phenotypes. To date, no connections have been found between these candidate genes and DCD-related etiology. Therefore, in the future, it will be of interest to determine if these molecules play a role in DCD phenotype regulation by performing genetic-association and functional studies that help to illuminate the etiology of DCD.

An important aspect of this work is the identification that different DCD-related phenotypes segregate in the BXD population – that is strains can be poor on some tasks, while being good on others. This allows us to genetically dissect different DCD-related phenotypes, and will allow us to identify genes underlying specific aspects of the disorder. This is important, as this can be used to inform findings from human studies of DCD – although human GWAS may be able to identify genes associated with DCD^87^, by combining with animal studies we will be able to identify which specific phenotypes within the disorder the gene contributes to.

The replication of these studies should be conducted in additional BXD strains to corroborate the preliminary QTL identified here, and to explore variation in motor coordination and corresponding QTLs. Subsequent papers of this study will address the variation in motor learning and corresponding QTLs and assess whether we can access phenotypes and genotypes relevant to DCD. Additionally, brain structural variability will be explored of BXD strains of mice to correlate the structural and behavioral differences seen in these 12 strains of mice.

## SUPPLEMENTARY FIGURES AND TABLE

**Figure S1.**
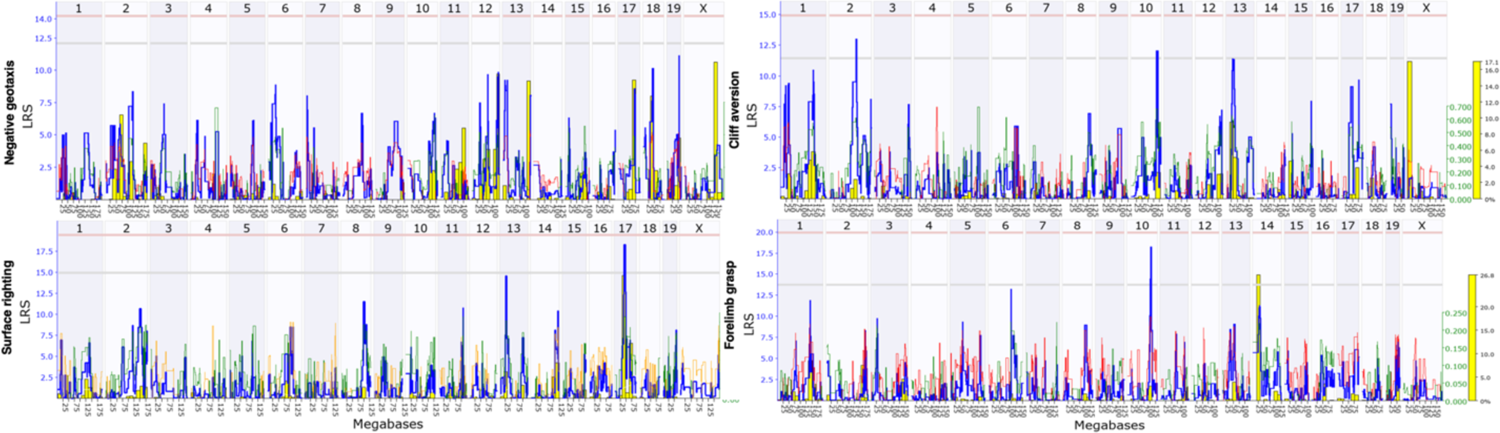
Genome-wide linkage map of negative geotaxis, surface righting, cliff aversion, and forelimb grasp on the fox neurodevelopmental battery to determine nervous system maturation. The overall blue trace shows the LRS. The genome-wide QTL map showing suggestive QTLs on Chromosome 17, 2, 10 and 13 for fox neurodevelopmental battery parameters. No QTL identified in negative geotaxis. The lower gray horizontal line represents suggestive LRS genome-wide threshold at p ≤ 0.63. The upper pink horizontal line represents significant LRS genome-wide threshold at p ≤ 0.05.

**Figure S2.**
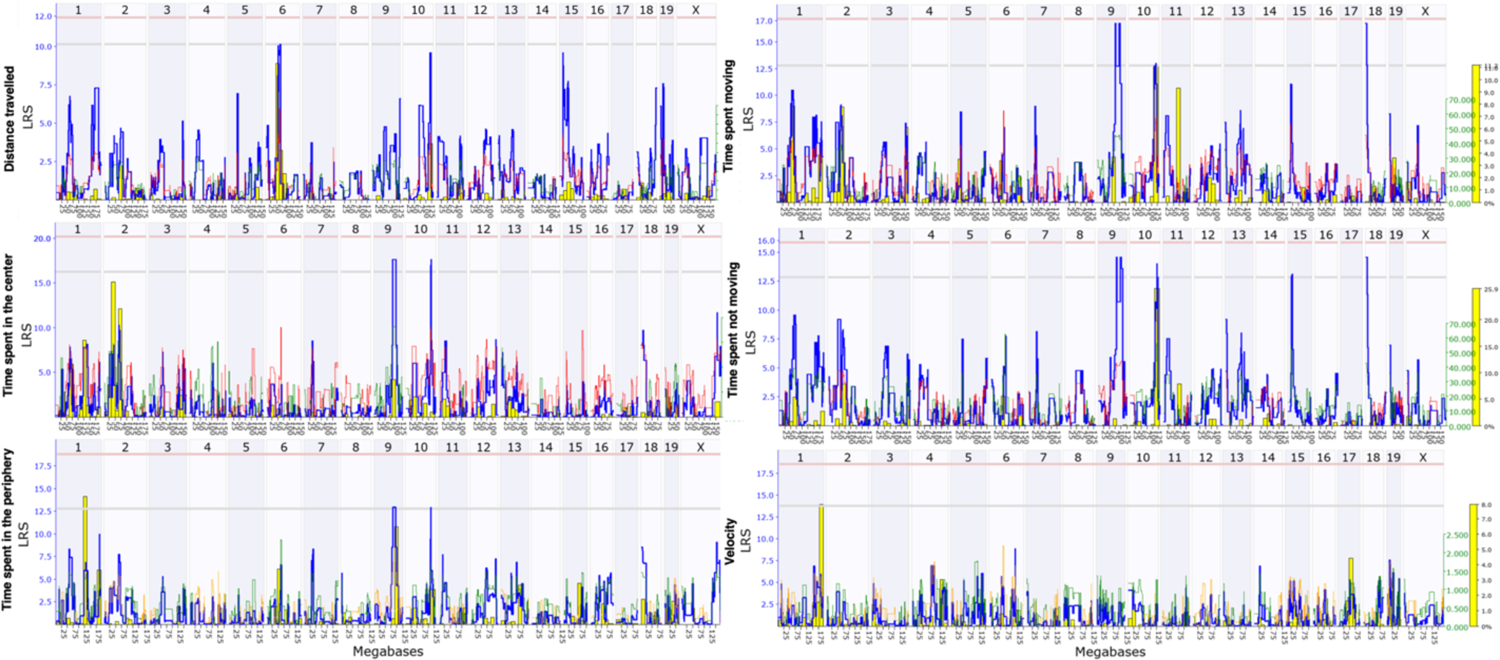
Genome-wide linkage map of distance travelled, time spent in the center, time spent in the periphery, time spent moving, time spent not moving and velocity of open field test to determine locomotor activity. The overall blue trace shows the LRS. The genome-wide QTL map showing suggestive QTLs on Chromosome 6, 9, 10, 15 and 18 on open field parameters. No QTL identified in velocity parameter. The lower gray horizontal line represents suggestive LRS genome-wide threshold at p ≤ 0.63. The upper pink horizontal line represents significant LRS genome-wide threshold at p ≤ 0.05.

**Figure S3.**
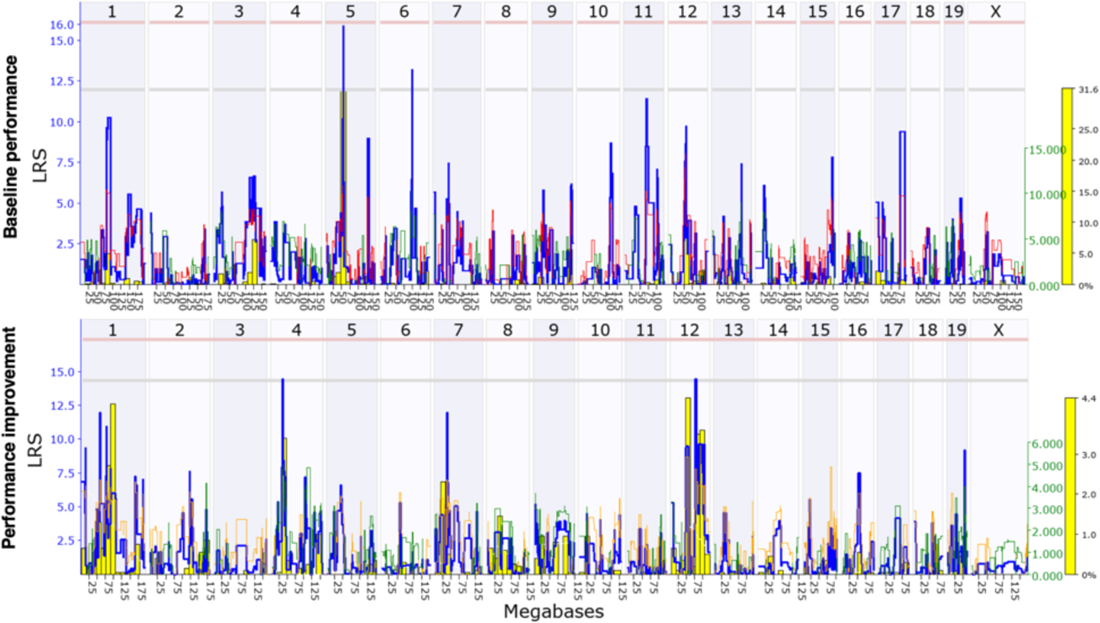
Genome-wide linkage map of baseline performance and improvements of standard rotarod to determine motor coordination and balance. The overall blue trace shows the LRS. The genome-wide QTL map showing suggestive QTLs on Chromosome 5, 6, 4 and 12 on standard rotarod. The lower gray horizontal line represents suggestive LRS genome-wide threshold at p ≤ 0.63. The upper pink horizontal line represents significant LRS genome-wide threshold at p ≤ 0.05.

**Figure S4.**
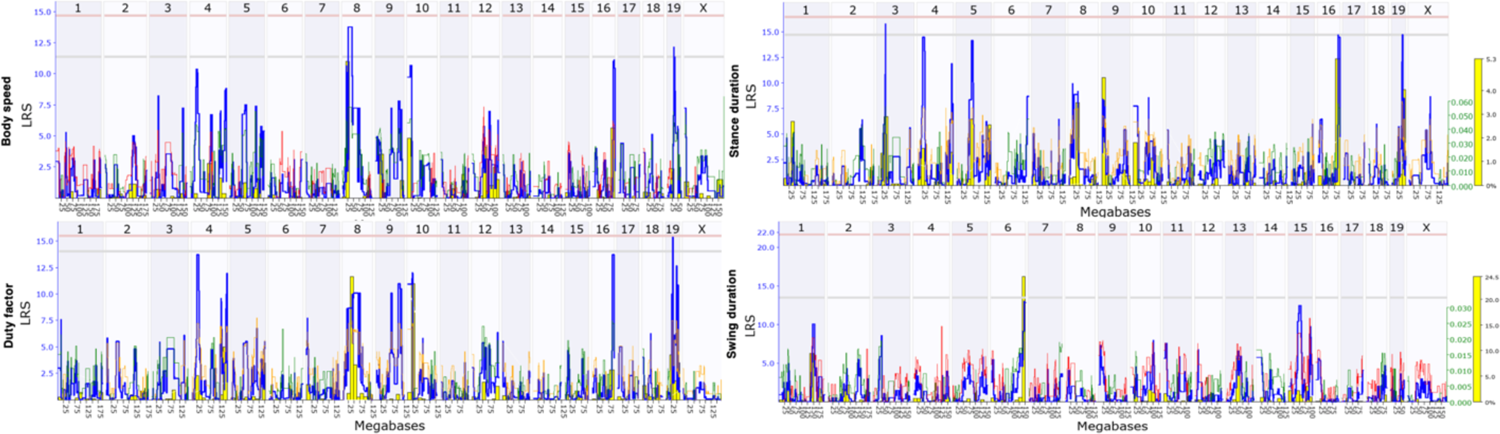
Genome-wide linkage map of body speed, duty factor, stance duration and swing duration of gait analysis to determine postural control. The overall blue trace shows the LRS. The genome-wide QTL map showing suggestive QTLs on Chromosome 5, 6, 4 and 12 on standard rotarod. The lower gray horizontal line represents suggestive LRS genome-wide threshold at p ≤ 0.63. The upper pink horizontal line represents significant LRS genome-wide threshold at p ≤ 0.05.

---

**Table S1.**
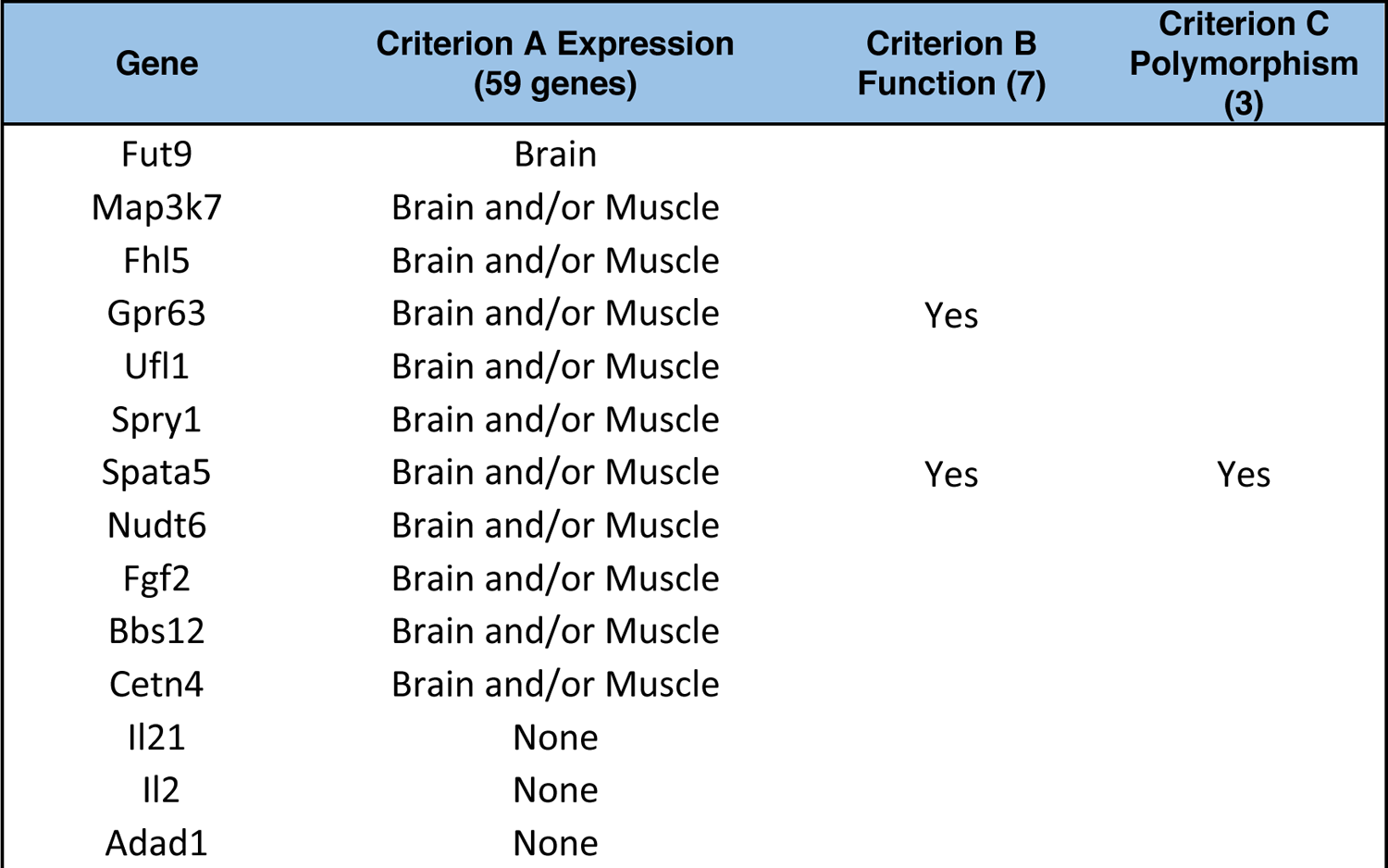
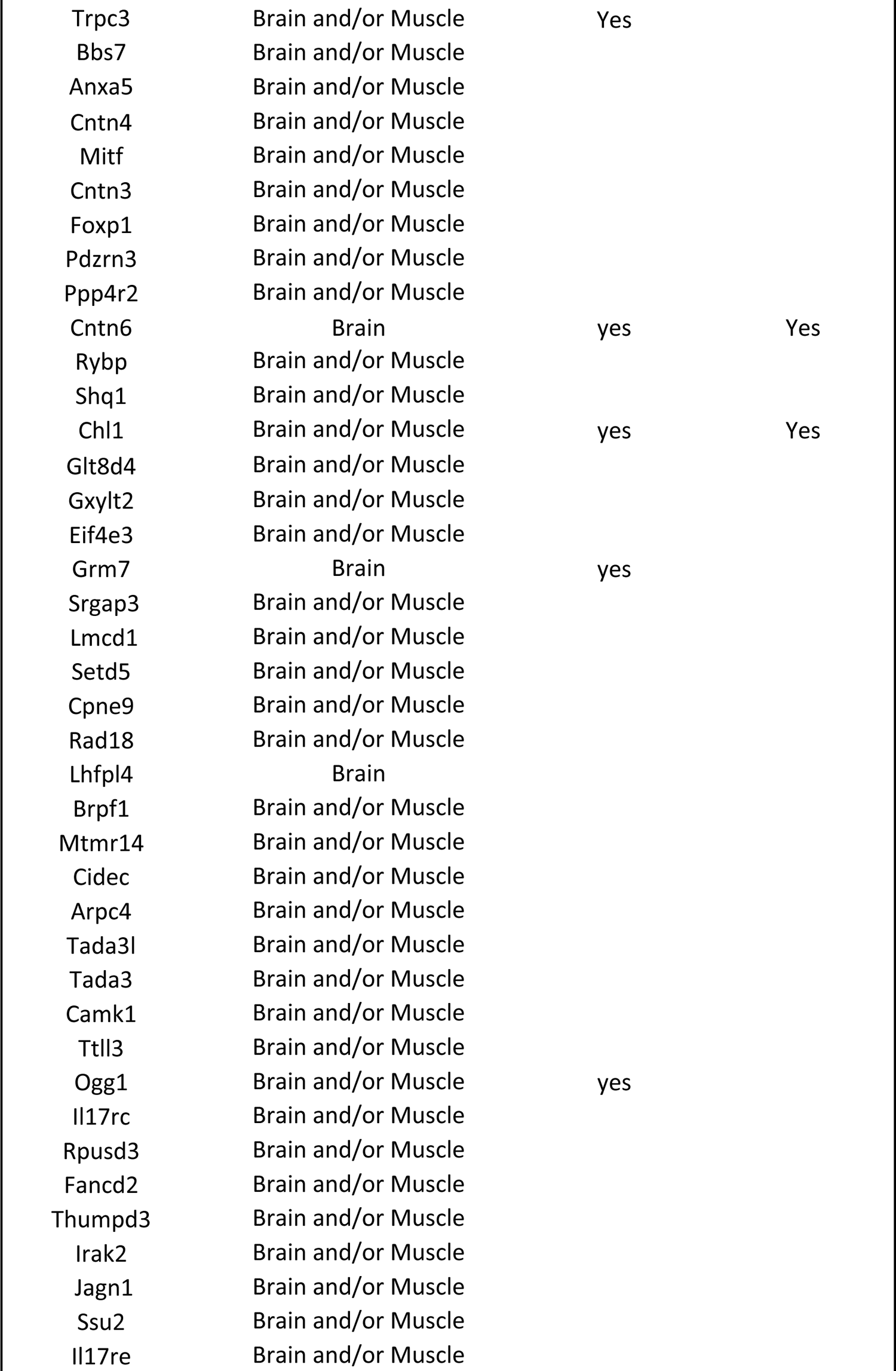
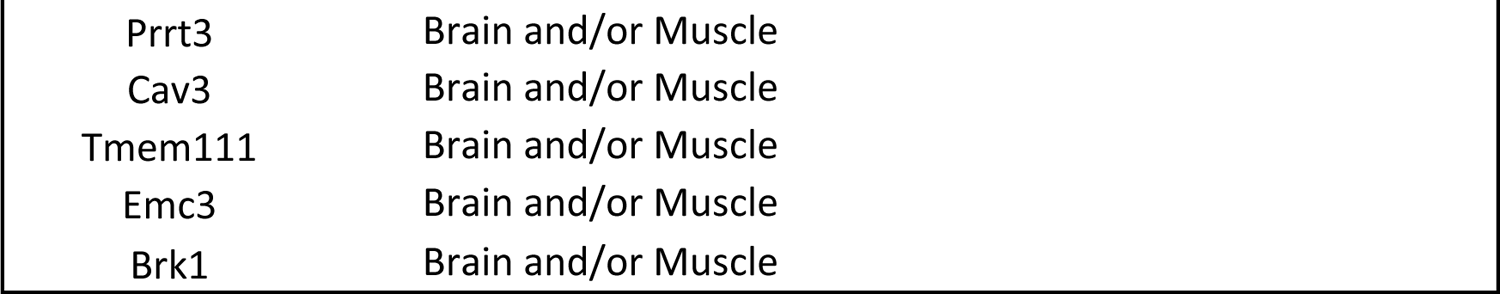
Identification of priority genes using three criterion. Empty areas show genes that did not meet the criterion system.

| Gene | Criterion A Expression<br>(59 genes) | Criterion B<br>Function (7) | Criterion C<br>Polymorphism<br>(3) |
| --- | --- | --- | --- |
| Fut9 | Brain |  |  |
| Map3k7 | Brain and/or Muscle |  |  |
| Fhl5 | Brain and/or Muscle |  |  |
| Gpr63 | Brain and/or Muscle | Yes |  |
| Ufl1 | Brain and/or Muscle |  |  |
| Spry1 | Brain and/or Muscle |  |  |
| Spata5 | Brain and/or Muscle | Yes | Yes |
| Nudt6 | Brain and/or Muscle |  |  |
| Fgf2 | Brain and/or Muscle |  |  |
| Bbs12 | Brain and/or Muscle |  |  |
| Cetn4 | Brain and/or Muscle |  |  |
| Il21 | None |  |  |
| Il2 | None |  |  |
| Adad1 | None |  |  |
| Trpc3 | Brain and/or Muscle | Yes |  |
| Bbs7 | Brain and/or Muscle |  |  |
| Anxa5 | Brain and/or Muscle |  |  |
| Cntn4 | Brain and/or Muscle |  |  |
| Mitf | Brain and/or Muscle |  |  |
| Cntn3 | Brain and/or Muscle |  |  |
| Foxp1 | Brain and/or Muscle |  |  |
| Pdzrn3 | Brain and/or Muscle |  |  |
| Ppp4r2 | Brain and/or Muscle |  |  |
| Cntn6 | Brain | yes | Yes |
| Rybp | Brain and/or Muscle |  |  |
| Shq1 | Brain and/or Muscle |  |  |
| Chl1 | Brain and/or Muscle | yes | Yes |
| Glt8d4 | Brain and/or Muscle |  |  |
| Gxylt2 | Brain and/or Muscle |  |  |
| Eif4e3 | Brain and/or Muscle |  |  |
| Grm7 | Brain | yes |  |
| Srgap3 | Brain and/or Muscle |  |  |
| Lmcd1 | Brain and/or Muscle |  |  |
| Setd5 | Brain and/or Muscle |  |  |
| Cpne9 | Brain and/or Muscle |  |  |
| Rad18 | Brain and/or Muscle |  |  |
| Lhfpl4 | Brain |  |  |
| Brpf1 | Brain and/or Muscle |  |  |
| Mtmr14 | Brain and/or Muscle |  |  |
| Cidec | Brain and/or Muscle |  |  |
| Arpc4 | Brain and/or Muscle |  |  |
| Tada3l | Brain and/or Muscle |  |  |
| Tada3 | Brain and/or Muscle |  |  |
| Camk1 | Brain and/or Muscle |  |  |
| Ttll3 | Brain and/or Muscle |  |  |
| Ogg1 | Brain and/or Muscle | yes |  |
| Il17rc | Brain and/or Muscle |  |  |
| Rpusd3 | Brain and/or Muscle |  |  |
| Fancd2 | Brain and/or Muscle |  |  |
| Thumpd3 | Brain and/or Muscle |  |  |
| Irak2 | Brain and/or Muscle |  |  |
| Jagn1 | Brain and/or Muscle |  |  |
| Ssu2 | Brain and/or Muscle |  |  |
| Il17re | Brain and/or Muscle |  |  |

|  |  |
| --- | --- |
| Prrt3 | Brain and/or Muscle |
| Cav3 | Brain and/or Muscle |
| Tmem111 | Brain and/or Muscle |
| Emc3 | Brain and/or Muscle |
| Brk1 | Brain and/or Muscle |

